# Exploration and activity show contrasting familial resemblance

**DOI:** 10.64898/2026.10.13.758810

**Authors:** Elisa Morbiato, Alexandra Glavaschi

**Affiliations:** Department of Biology, University of Padova, 35121 Padova, Italy; Institute for Vertebrate Biology, Czech Academy of Sciences, Brno, Czech Republic

**Keywords:** Bayesian modelling, inclusive inheritance, animal personality, behavioural modularity, phenotypic integration

## Abstract

Whether complex behaviours that appear unitary are governed by a single integrated system or composed of distinct evolutionary modules remains unclear. For instance, an animal’s locomotion trajectory depends on two independent decisions: how much to move versus where to go. Determining whether these behavioural dimensions are linked or vary independently is essential, as modular traits impact the evolutionary potential of complex phenotypes. We investigated this framework in newborn Trinidadian guppies (*Poecilia reticulata*) by decoupling a single locomotor sequence into its exploratory and activity components and evaluating their structure of inclusive inheritance through a nested paternal half-sib breeding design. Our findings reveal that these behavioural dimensions exhibit contrasting patterns of familial resemblance, with exploration being structured with fathers, whereas activity with mothers. These findings indicate potential evolutionary decoupling between how much an animal moves and where it goes, supported by trait-specific structures of inclusive inheritance. Importantly, this modular architecture suggests that different components of a single locomotor sequence can be targeted independently by natural selection, providing a powerful mechanism to bypass rigid behavioural syndromes and drive the diversification of complex personality profiles.

## Introduction

Behaviours that appear unitary are often composed of partially separable dimensions [1]. For instance, foraging depends both on the effort invested in searching and on where that effort is allocated; dispersal involves both the decision to leave and the distance travelled. Also, courtship can vary independently in its rate, intensity and structural complexity [2,3]. These components may be coordinated because they contribute to a common behavioural function, but functional coordination does not necessarily imply a shared underlying regulatory structure. Different components of behaviour may indeed be governed by distinct genetic, developmental, neural, or physiological mechanisms [4,5]. Because these regulatory mechanisms can operate independently, the resulting patterns of phenotypic variability among individuals may differ significantly from one behavioural component to another. Phenotypic integration describes the tendency of multiple components to covary and function as a coordinated unit. Conversely, modularity refers to the independent variation of distinct dimensions, which allows them to vary and evolve semi-autonomously [5–7]. Assessing whether behaviours extracted from a single sequence are integrated or modular is crucial, as aggregate measures can obscure independent pathways of evolutionary potential [8].

Locomotion provides a powerful test of this framework. An animal’s trajectory is indeed dictated by two fundamental behavioural decisions that address distinct ecological questions: *how much to move* (hereafter activity) and *where to go* (hereafter exploration) which represent two core axes of animal personality [3]. Crucially, although these decisions generate different motor outputs through a single, functionally unitary behaviour, how an individual balances them shapes the organization of behavioural syndromes [9]. For instance, one individual may move continuously while remaining within a restricted, safe area, whereas another may enter exposed or unfamiliar space despite moving only briefly. Resolving these two behavioural decisions in different ways gives rise to distinct personality profiles, with the extreme ends characterized by a highly active but risk-averse phenotype versus a low-activity, risk-tolerant one. If these outputs are strictly integrated, the resulting syndrome is evolutionarily constrained; conversely, if individuals can decouple *how much they move* from *where they go*, the combinations of behavioural phenotypes increase, boosting the evolutionary potential. Nevertheless, distinct metrics extracted simultaneously from a single locomotor sequence are often treated as interchangeable indicators of an underlying behavioural tendency or combined into composite scores [8,10,11]. This practice implicitly assumes that activity and exploration are not only phenotypically integrated but also share an underlying regulatory structure.

One way of testing whether these behavioural components are integrated or modular is examining how variation is transmitted across generations. Because traits can be transmitted through different mechanisms (such as genes, non-genetic parental effects, and the environment) [12–14], studying the structure of familial resemblance, namely how variance is partitioned between mothers and fathers, allows us to map this inclusive inheritance [15] and determine whether behaviours are integrated or modular. Importantly, these parental and environmental sources of variation need not contribute equally to all components of behaviour. In guppies, genetic and maternal contributions differ among behaviours measured in open-field assays [16], while studies in rodents show that maternal genetic and non-genetic effects can differentially influence exploration and anxiety-like behaviour [17]. Paternal resemblance may likewise arise through independent genetic inheritance or non-genetic paternal effects transmitted before or after fertilization [18]. Whether components extracted from within a single behavioural sequence are evolutionarily integrated or modular, however, remains largely unresolved.

To resolve how these behavioural components are structured, we evaluated two alternative hypotheses regarding the structuring of familial resemblance in locomotion. Under an integrated phenotype hypothesis, activity and exploration represent coupled dimensions of a single behavioural trait. We therefore predicted that they would exhibit strong phenotypic covariation and a shared pattern of familial variance allocation. Under a modular phenotype hypothesis [1,5], activity and exploration represent distinct units. We consequently predicted weak phenotypic covariation and trait-specific structuring of familial resemblance, with variance in each behaviour being explained by different parental sources.

We tested these predictions in newborn Trinidadian guppies (*Poecilia reticulata*) using an open-field assay and a nested paternal half-sib breeding design that separated sire-associated resemblance from additional resemblance among offspring sharing the same dam. Studying offspring within three days of birth allowed us to examine familial structuring near the beginning of behavioural development, before individuals had accumulated divergent postnatal experiences [19]. We quantified *activity* as the proportion of time spent mobile and *exploration* as the proportion of time spent in the exposed central region of the arena. Using a Bayesian bivariate hierarchical model, we partitioned variation in each behaviour into sire-associated, dam-within-sire-associated and residual components. Given that these components may combine genetic, parental and shared-environmental influences, we interpret them as sources of familial resemblance driven by inclusive inheritance, instead of narrow-sense heritability.

## Materials & Methods

### Study population and breeding design

Experimental guppies (*Poecilia reticulata*) originated from a semi-natural population established at the Botanical Garden of Padova in 2014 using fish descended from a population collected from the Lower Tacarigua River, Trinidad, in 2003. The laboratory population was maintained at the University of Padova in 150-litre stock aquaria at a density of approximately one fish per litre, under a 12:12 light:dark photoperiod and at a water temperature of 26° C ± 1. Juveniles were collected regularly from the stock population and separated by sex at sexual maturation to obtain virgin females and males of known age. All fish were fed ad libitum with dry food and *Artemia sp*. nauplii twice daily.

To establish the experimental families and generate the paternal half-sibships, adult males aged 4– 6 months were sequentially paired with three virgin females on consecutive days, remaining with each female for 24 hours. This design produced paternal half-sib families where offspring from the same male but different females shared a paternal genetic line but developed under distinct maternal environments. Following mating, females were housed individually in 3.5-litre aquaria fitted with nursery compartments. Tanks were inspected twice daily until parturition or for a maximum of 60 days.

**Figure 1.**
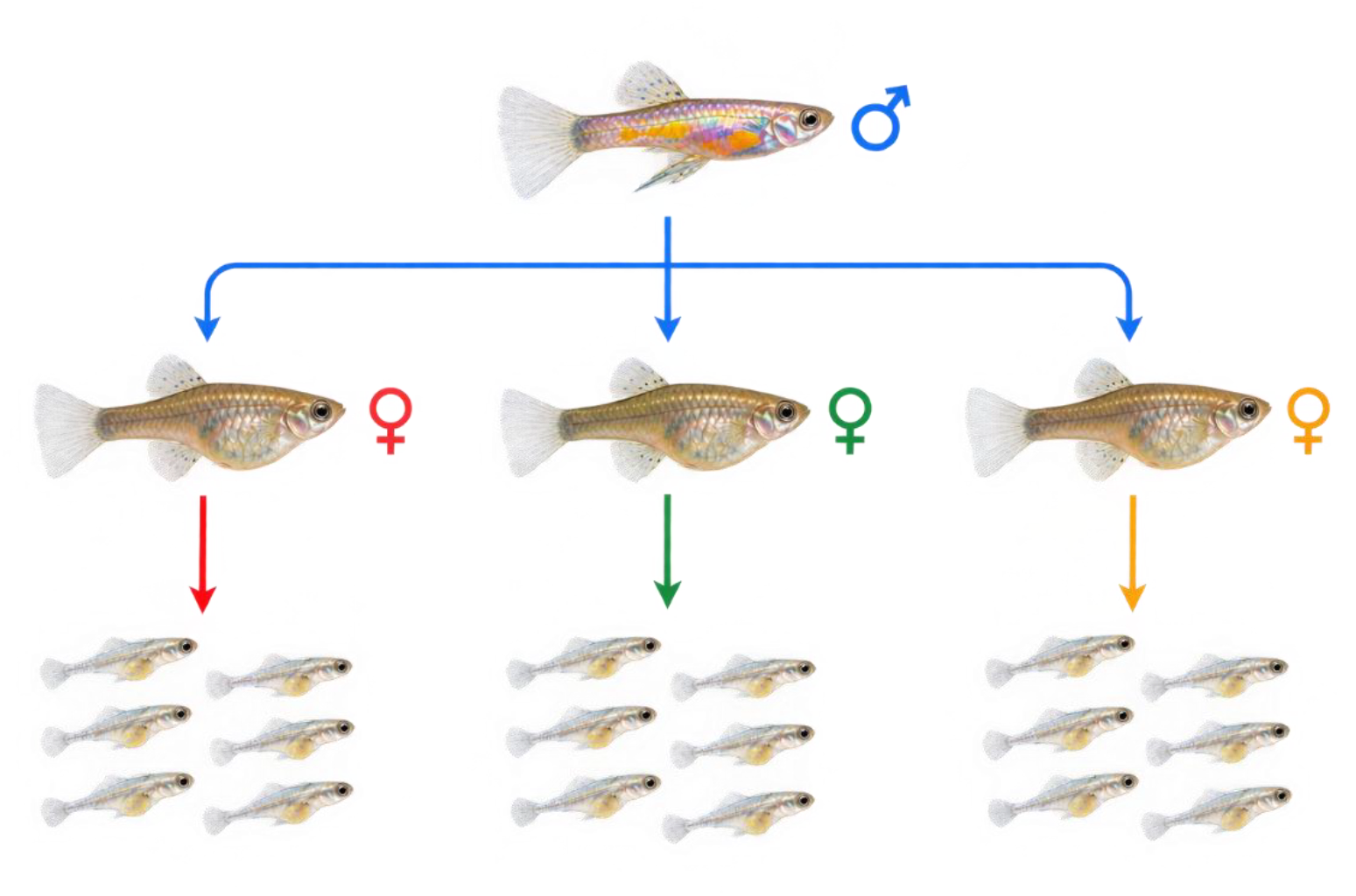
Nested paternal half-sib breeding design layout. Individual adult males (sire line, in blue) were sequentially paired with three virgin females (dams in red, green, and yellow) for 24-hour periods. Offspring within the same maternal brood share both a paternal and maternal genetic line and a common gestational environment (full-sibs). Conversely, offspring across different broods (e.g., red versus green) share the same paternal genes but different maternal genes and environment (half-sib).

### Behavioural assay

Within three days of birth, each offspring was individually subjected to an open-field test adapting Breedveld et al. [20]. Briefly, each fish was gently transferred to a circular white arena 18 cm in diameter and containing 300 mL of water maintained at 26 ± 1°C. The arena was video recorded from above with a Panasonic HCV180 camera. Fish were allowed to acclimate for 5 min and were subsequently filmed for 10 min while freely exploring the arena in the absence of external disturbance. The arena was washed with deionized water, rinsed, and dried and the water replaced between trials to minimise chemical cues from previously tested fish.

The behavioural dataset contained open-field trial observations from 232 offspring from 56 dams and 35 sires. One offspring lacked a valid tracking measure, resulting in 231 individuals with complete measurements for the bivariate analysis. The intended paternal half-sib design was reduced by female mortality and reproductive failure, resulting in an unbalanced family structure. Eighteen sires produced offspring with one dam, thirteen with two dams, and four with three dams. The number of offspring per sire ranged from 2 to 21, with a median of 6.

### Behavioural tracking

Fish trajectories were extracted from the recordings using ToxTrac [21] at a sampling rate of 25 frames s^-1^. Within ToxTrac, two nested spatial regions were defined digitally: Zone 1 encompassed the entire arena, whereas Zone 2 consisted of a central circle measuring 12 cm in diameter. The dimension of the inner zone was defined following previous studies that investigated exploratory behaviour and tendencies to thigmotaxis in guppies [10]. Time spent in Zone 2 represented time in the central, more exposed portion of the arena and was used as the proxy of exploration, calculated by dividing the time in Zone 2 by the effective tracking duration. Activity was quantified using the mobility rate exported by ToxTrac, defined as the proportion of the effective tracking period during which an individual was classified as mobile (i.e. speed exceeding 1 mm s^−1^). The phenotypic association between behavioural endpoints was described using Spearman’s rank correlation.

### Statistical analyses

All analyses were conducted in R version 4.4.2 (R Core Team 2024) using *brms* 2.23.0 [22] and *posterior* 1.6.1.

### Bivariate hierarchical model

Exploration and activity were analysed as the two primary behavioural endpoints. Because both variables were expressed as proportions bounded between zero and one, they were logit-transformed to meet the assumptions of Gaussian modelling. A standard continuity correction (ε = 0.5/600) was applied to boundary observations prior to transformation to avoid undefined logits. Finally, each transformed endpoint was standardised (mean = 0, SD = 1) to place both responses on a common scale and allow the use of identical informative priors.

We fitted a Bayesian bivariate Gaussian hierarchical model in which activity and exploration were analysed jointly. Each behavioural response was modelled using an intercept, a random intercept for sire and a random intercept for dam nested within sire. The bivariate formulation estimated separate sire-associated, dam-within-sire-associated and residual variances for each endpoint. It also estimated correlations between exploration and activity at the sire, dam-within-sire and residual levels. For each endpoint and posterior draw, sire-associated and dam-within-sire-associated variance partition coefficients were calculated by dividing the corresponding variance component by the sum of sire-associated, dam-within-sire-associated and residual variance.

Our inferential question was whether the ordering of sire-associated and dam-within-sire-associated variance differed between the two behavioural endpoints. For exploration, we calculated the sire-associated minus dam-within-sire-associated variance partition coefficient (VPC) difference; for activity, we calculated the dam-within-sire-associated minus sire-associated VPC difference. Positive values therefore indicated the predicted ordering within each endpoint. We quantified joint posterior support for the opposing ordering as the proportion of posterior draws in which sire-associated variance exceeded dam-within-sire-associated variance for exploration and dam-within-sire-associated variance exceeded sire-associated variance for activity, with both conditions occurring simultaneously. We also calculated a between-trait log variance-ratio contrast, defined as the log sire-to-dam variance ratio for exploration minus the corresponding log ratio for activity. Positive values indicated that the sire-to-dam variance ratio was greater for exploration than for activity. This contrast quantified the magnitude of the relative difference between endpoints, whereas evidence for the opposing ordering was provided by the within-endpoint comparisons and their joint posterior probability. Posterior distributions were summarised using medians, 95% credible intervals and posterior probabilities that the planned contrasts were in the predicted direction.

### Priors and model estimation

Weakly informative priors were used throughout. Intercepts were assigned normal priors with a mean of zero and a standard deviation of 0.5. Familial standard deviations were assigned Student-t priors with three degrees of freedom and a scale of 0.5, while residual standard deviations used the same distribution with a scale of 0.7. Correlation matrices were assigned LKJ (2) priors, providing mild regularisation towards weaker correlations. The primary model was estimated using four Markov chains of 6,000 iterations, including 3,000 warm-up iterations per chain. We used an adapt delta of 0.995 and a maximum tree depth of 14. Posterior predictive checks were used to evaluate whether datasets generated by the fitted model reproduced the principal distributional characteristics of both behavioural endpoints.

### Sensitivity and influence analyses

We assessed sensitivity to prior specification by refitting the model with wider priors for intercepts and variance components and LKJ (1) priors for the correlation matrices. Because sires represented by a single dam provide no direct within-sire replication for separating sire-associated from dam-within-sire-associated variance, we also repeated the analysis using only sires represented by at least two dams. We additionally conducted a leave-one-sire-out influence analysis. The model was refitted after excluding each sire in turn, and the between-trait log variance-ratio contrast was recalculated for every reduced dataset. This analysis assessed whether the principal pattern depended disproportionately on a single paternal family. Each reduced model was evaluated using the same convergence and sampling diagnostics as the primary model. Convergence and sampling performance were assessed using rank-normalised 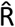, bulk and tail effective sample sizes, divergent transitions and maximum-tree-depth diagnostics.

## Results

### Familial variance in activity and exploration

Exploration and activity were weakly correlated at the phenotypic level (Spearman’s ρ = − 0.09).

Familial variance differed across the two behavioural dimensions (Figure 1). For exploration, the posterior median sire-associated variance partition coefficient was 0.392 (95% credible interval, CrI: 0.163–0.579), whereas the dam-within-sire-associated coefficient was 0.047 (95% CrI: 0.000–0.266). The posterior median difference between these components was 0.343 (95% CrI: − 0.086–0.56), with a posterior probability of 0.956 that the sire-associated component was larger.

**Figure 1a.**
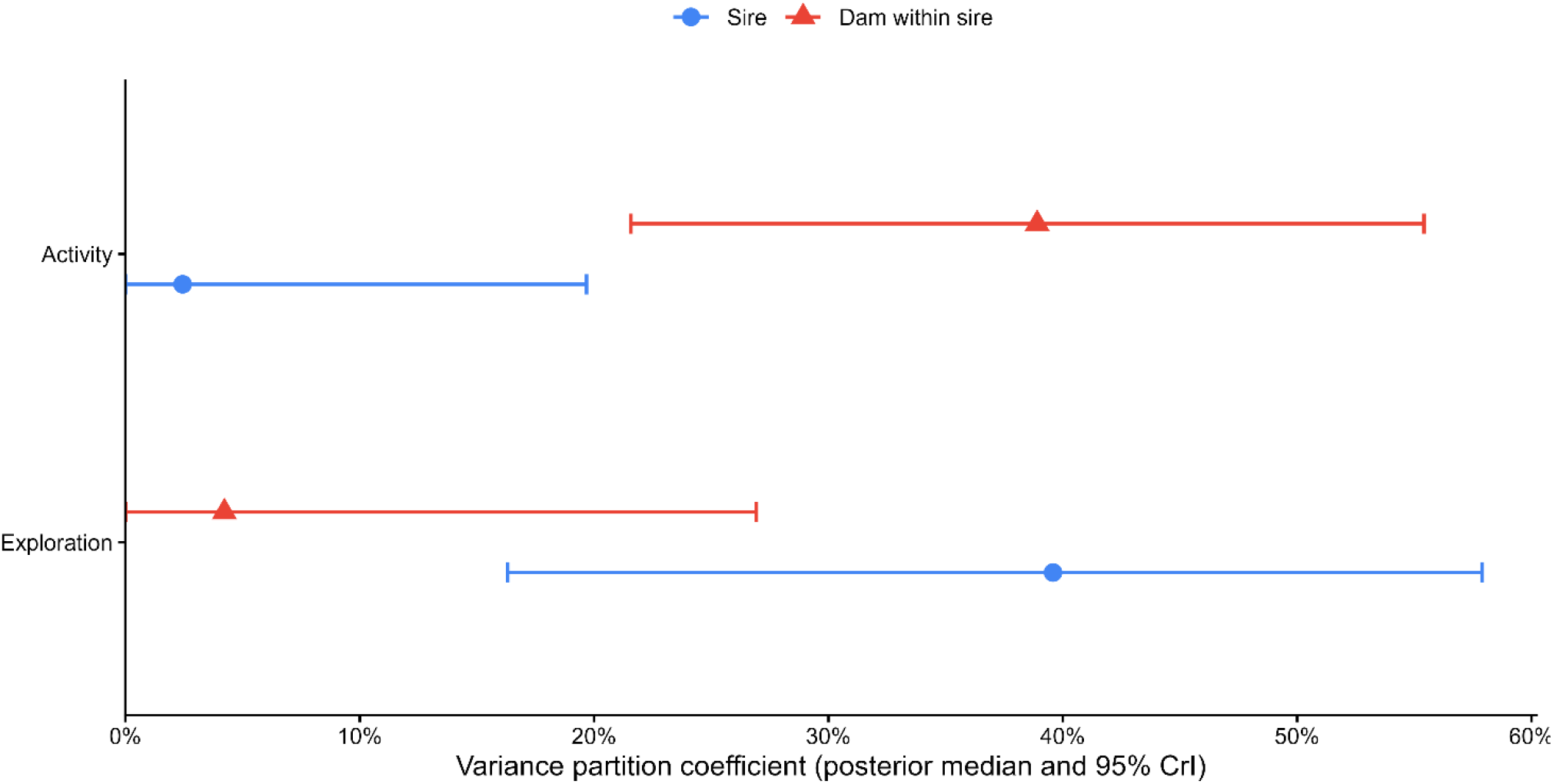
Familial variance partitioning of activity and exploration. Points show posterior median variance partition coefficients (VPC) and horizontal bars show 95% credible intervals for sire-associated variance (blue circles) and dam-within-sire-associated variance (red triangles). For exploration, the sire-associated coefficient was 0.392 (95% CrI 0.163– 0.579), compared with 0.047 (0.000–0.266) for dams within sires. Activity showed the reverse pattern, with coefficients of 0.024 (0.000–0.197) for sires and 0.389 (0.216–0.554) for dams within sires. Exploration was calculated as the proportion of effective tracking time spent in the central zone and activity as the proportion of time classified as mobile.

Activity showed the opposite pattern. Its posterior median sire-associated variance partition coefficient was 0.024 (95% CrI: 0.000–0.197), compared with 0.389 (95% CrI: 0.216–0.554) for the dam-within-sire-associated component. The posterior median difference in the predicted direction was 0.358 (95% CrI: 0.049–0.543), with a posterior probability of 0.986 that the dam-within-sire-associated component was larger. The predicted opposing ordering occurred simultaneously for both endpoints in 0.943 of the posterior draws.

### Trait-specific contrast in familial resemblance

The opposing variance patterns produced a pronounced difference in familial structure between exploration and activity (Figure 2). The posterior median between-trait log variance-ratio contrast was 5.325 (95% CrI: 1.142–13.321). The posterior probability that this contrast was positive was 0.993, providing strong support for a greater sire-to-dam variance ratio in exploration than in activity.

**Figure 2.**
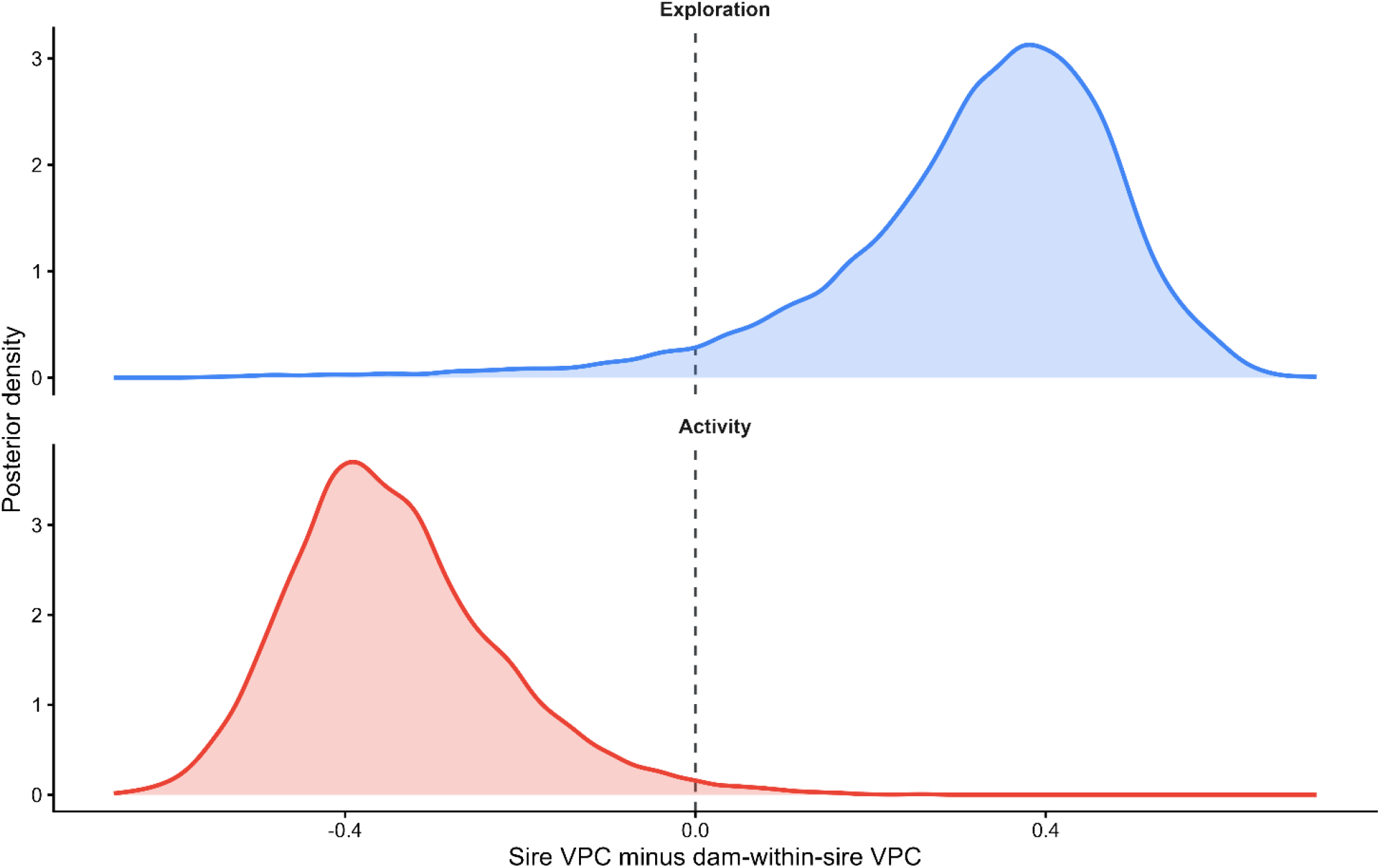
Posterior contrasts between sire-associated and dam-within-sire-associated variance. Curves show the posterior distributions of sire-minus-dam-within-sire variance partition coefficient (VPC) differences for exploration and activity. For exploration, positive values indicate a larger sire-associated VPC; the posterior median difference was 0.343 (95% CrI: −0.086 to 0.560; P_post_ = 0.957). For activity, negative values indicate a larger dam-within-sire-associated VPC; the posterior median difference was −0.358 (95% CrI: −0.543 to −0.049; P_post_ = 0.986). The overall crossover contrast comparing sire-to-dam variance ratios between endpoints was 5.325 (95% CrI: 1.142 to 13.321; P_post_ = 0.993).

### Sensitivity and influence analyses

Sampling diagnostics for the primary model were satisfactory, with no divergent transitions or maximum-tree-depth exceedances. Posterior predictive checks reproduced the overall location and dispersion of both behavioural endpoints. The between-trait log variance-ratio contrast remained positive under weaker priors, indicating that the inferred between-trait difference in familial variance structure was not driven by prior regularisation. It also remained positive when the analysis was restricted to the 17 sires represented by at least two dams. This restriction excluded the 18 sires represented by a single dam, for which sire-associated and dam-within-sire-associated variation are less readily distinguished, and therefore confirmed that the opposing pattern of familial variance components was supported by the replicated part of the family design (electronic supplementary material, Table S2).

The leave-one-sire-out analysis provided a complementary assessment of influence. The posterior median contrast remained positive after each of the 35 sires was excluded in turn (electronic supplementary material, Table S3 and Figure S1). Posterior support was lowest after the exclusion of sire m58 but remained high (P_post_ [C > 0] = 0.973).

Thus, the opposing ordering of familial variance components was robust to prior specification, was retained among sires with replication across dams, and was not driven by any single paternal family.

## Discussion

Behaviours expressed simultaneously through the same locomotor sequence are often assumed to reflect a single, integrated axis of individual variation [3,9]. Our results challenge this view. Activity and exploration were weakly correlated, and both exhibited familial resemblance structured at opposing hierarchical levels: exploration was structured to sire identity, whereas activity was structured among dams within sires. This contrast indicates that the duration of movement and its spatial allocation are not integrated. Instead, they represent distinct evolutionary modules inherited by different familial sources.

Although our breeding design cannot disentangle narrow-sense genetic variance from non-genetic parental or environmental effects, our findings reveal that variation in these two modules is driven by two different parental sources. To our knowledge, this provides the first evidence that distinct components of a single, functionally unitary behavioural sequence can be inherited through different parental lines, offering a powerful new perspective on how behavioural modularity evolves.

The inheritance of modularity can be achieved by genetic and non-genetic mechanisms where maternal and paternal lines target separate components of behaviour independently [23]. For instance, maternal and early developmental environments can influence offspring traits through prenatal provisioning, hormones, or gestational stress, but these influences need not affect all behavioural dimensions equally [23]. In lesser black-backed gulls prenatal and postnatal social environments shape early exploratory activity but have no effect on anxiety-like responses [24]. Similarly, cross-fostering experiments in zebra finches demonstrate that offspring exploration resemble those of their foster parents rather than their genetic parents, highlighting a purely environmental pathway of transmission [25]. In guppies, maternal perception of predation risk increased offspring exploration, but had no effect on schooling behaviour, demonstrating that maternal effects can be behaviour-specific [26]. The mechanism underlying the prominent dam-within-sire-associated variance detected for activity cannot be distinguished in our design: it may encompass both genetic and non-genetic maternal effects. Nevertheless, previous work on juvenile guppies found substantial maternal effects on activity, but little evidence for additive genetic variance. [16]. The substantial sire-associated variance in exploration could reflect genetic inheritance or non-genetic effects transmitted through the ejaculate. Ejaculate-mediated paternal effects have been demonstrated experimentally in guppies, although their contribution to behavioural variation remains unknown [27].

Such behavioural specificity in parental influences provides strong empirical support for the modular phenotype hypothesis in other vertebrates, challenging the view that behavioural syndromes are always rigidly integrated. Crucially, our study demonstrates that two simultaneously expressed components of a single locomotion sequence—activity (*how much to move*) and exploration (*where to go*)—harbour decoupled structures of familial resemblance. This decoupling provides a clear evolutionary mechanism through which populations can break away from rigid behavioural syndromes to express alternative, flexible personalities.

Such modular systems of familial resemblance may lie at the origin of the vast diversity observed in individual behaviour. Behavioural modularity fundamentally alters both the direction and predictability of evolutionary change. If the quantity of movement and its spatial allocation are governed by independent familial processes, selection acting on one component will not generate a correlated response in the other. Rather than constituting a single, uniform personality axis, these independent behavioural modules can be recombined into multiple different permutations. This variational freedom allows natural selection to fine-tune specific ecological responses independently, acting as a driver for the radiation of diverse individual behavioural strategies. A population could therefore evolve greater use of exposed or unfamiliar space without a corresponding increase in general activity, or experience developmental changes in locomotor activity without altering spatial decisions. More generally, the evolutionary response of a complex behaviour cannot be predicted reliably from its phenotypic organisation alone but depends on how genetic, parental and environmental variance is distributed among its constituent components [4].

Our results show that a single locomotor trajectory can contain distinct modules with contrasting familial resemblance. More generally, decomposing behavioural sequences into their distinct variational modules can reveal discrete pathways of evolutionary potential that aggregate measures completely conceal. This distinction is critical both for mapping the true diversity of behavioural variation and for predicting how complex traits respond to selection.

### Ethics permit

This research was approved by the University of Padua’s Animal Ethics Committee (no. 256 /2018)

### Data accessibility

DOI: 10.25430/researchdata.cab.unipd.it.00001938

## Authors’ contributions

E.M. designed the study. All authors contributed towards data collection.

E.M. analysed the data and wrote the first draft of the paper. All authors contributed towards subsequent drafts and approved the final version of the manuscript.

## Acknowledgements

This study is part of a broader research project investigating the genetic basis of behaviour and life-history traits in male guppies conceived by Andrea Pilastro. We thank him for scientific, logistic, and financial support and for providing the guidance and the resources necessary to conduct this study. We are grateful to Andrea Pilastro and Ingo Schlupp for reviewing an earlier draft of the manuscript and providing valuable suggestions that improved the paper. We thank Andrea Bontorin for his contribution to the collection of behavioural data.

## Supplementary information

### Model diagnostics and posterior predictive assessment

The primary bivariate model showed satisfactory sampling performance (Table S1). The maximum rank-normalised 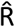 was 1.007, and the minimum bulk and tail effective sample sizes were 643 and 1111, respectively. No divergent transitions or maximum-tree-depth exceedances occurred.

**Table S1.**
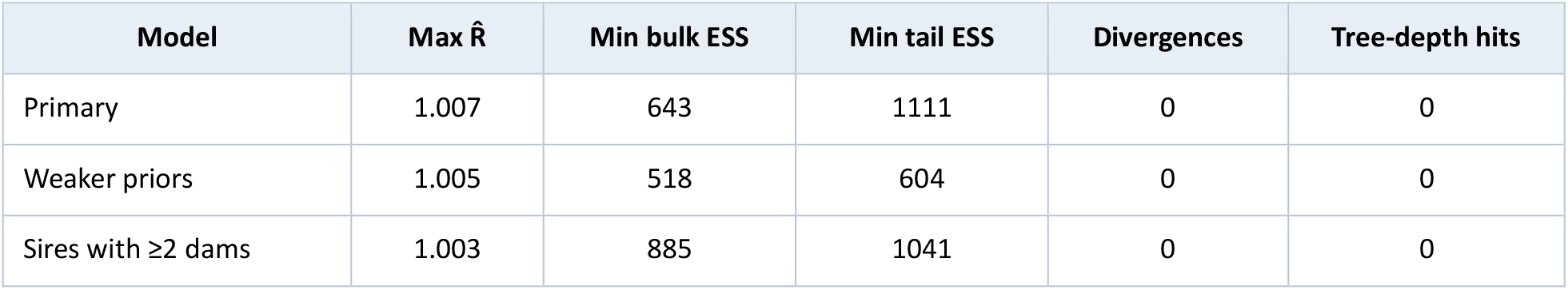
Sampling diagnostics for the primary and sensitivity models.

| Model | Max $\hat{R}$ | Min bulk ESS | Min tail ESS | Divergences | Tree-depth hits |
| --- | --- | --- | --- | --- | --- |
| Primary | 1.007 | 643 | 1111 | 0 | 0 |
| Weaker priors | 1.005 | 518 | 604 | 0 | 0 |
| Sires with $\geq 2$ dams | 1.003 | 885 | 1041 | 0 | 0 |

### Sensitivity to prior specification and family structure

The between-trait log variance-ratio contrast was consistent under both alternative prior specification and when the analysis was restricted to sires represented by at least two dams (Table S2). Under weaker priors, the posterior median contrast was 5.54 (95% credible interval, CrI: 1.24 – 13.30), and its posterior probability of being positive was 0.993. Restricting the analysis to sires represented by at least two dams produced a posterior median contrast of 5.14 (95% CrI: 0.86 – 12.73; posterior probability positive = 0.989). These estimates were similar to the primary model estimate of 5.32 (95% CrI: 1.14 – 13.32; posterior probability positive = 0.993). Sampling diagnostics were satisfactory for both sensitivity models (Table S1). Maximum rank-normalised 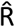 values were 1.005 under weaker priors and 1.003 in the analysis restricted to sires with at least two dams. Neither model produced divergent transitions or maximum-tree-depth exceedances.

**Table S2.**
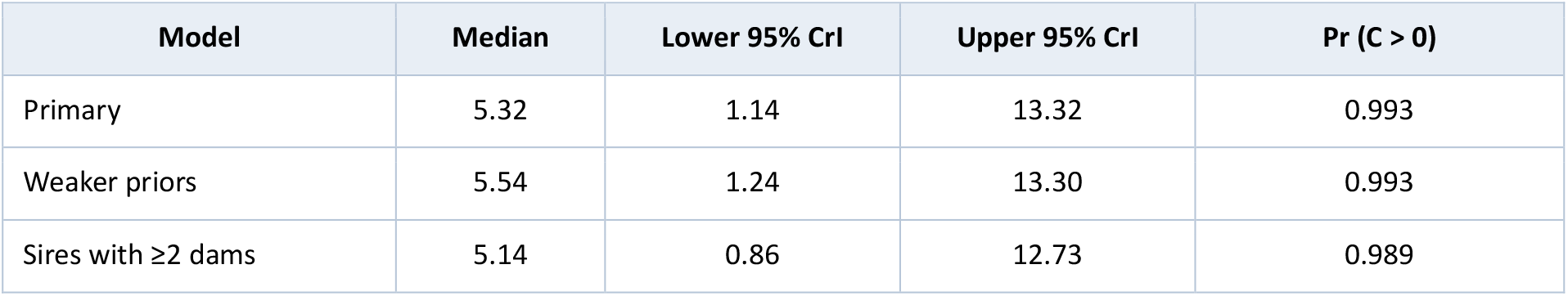
Between-trait log variance-ratio contrast under the primary and sensitivity models. Positive values indicate a greater sire-to-dam variance ratio for exploration than for activity.

### Leave-one-sire-out influence analysis

Excluding each of the 35 sires in turn did not alter the direction of the between-trait log variance-ratio contrast (Figure S1; Table S3). Posterior median contrast estimates ranged from 4.29 to 6.35. The minimum posterior probability that the contrast was positive was 0.973. The most conservative estimate followed exclusion of sire m58 (posterior median = 4.29, 95% CrI: − 0.09 – 12.16; posterior probability positive = 0.973). Excluding sire m2 produced the next-lowest median estimate (4.67, 95% CrI: 0.53 – 12.50; posterior probability positive = 0.988). Thus, the positive between-trait contrast was not attributable to a single paternal family.

Ten reduced models had a maximum parameter-level 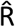 marginally above 1.01 (Table S4). In eight cases, the maximum value occurred for the dam-within-sire-level correlation between exploration and activity, in the remaining two, it occurred for the dam-within-sire standard deviation for exploration. Convergence diagnostics for the derived contrast were satisfactory in all ten models: 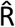 ranged from 1.000 to 1.003, bulk effective sample sizes from 810 to 1293, and tail effective sample sizes from 945 to 2508. No reduced model contained divergent transitions.

**Table S3.**
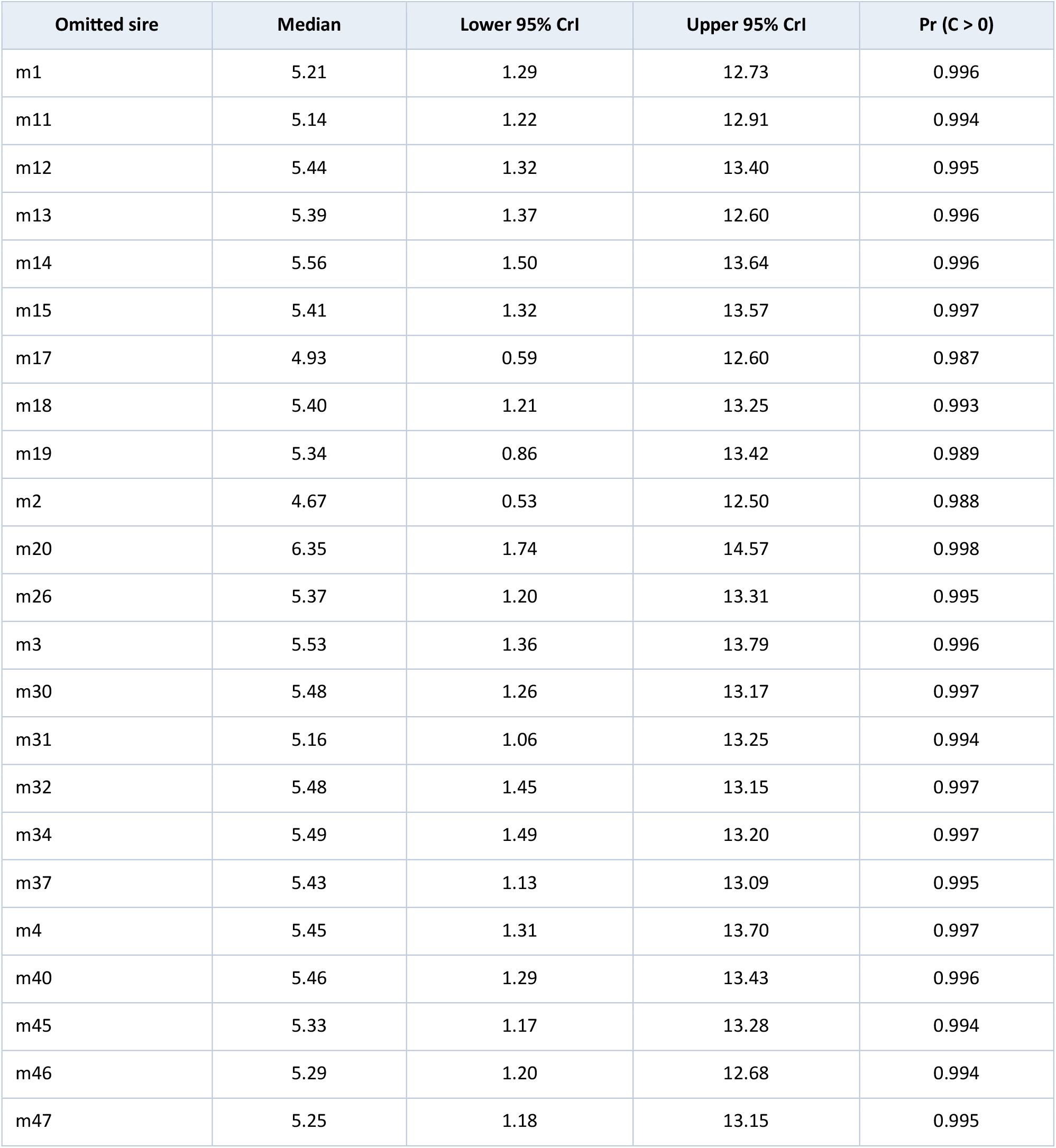
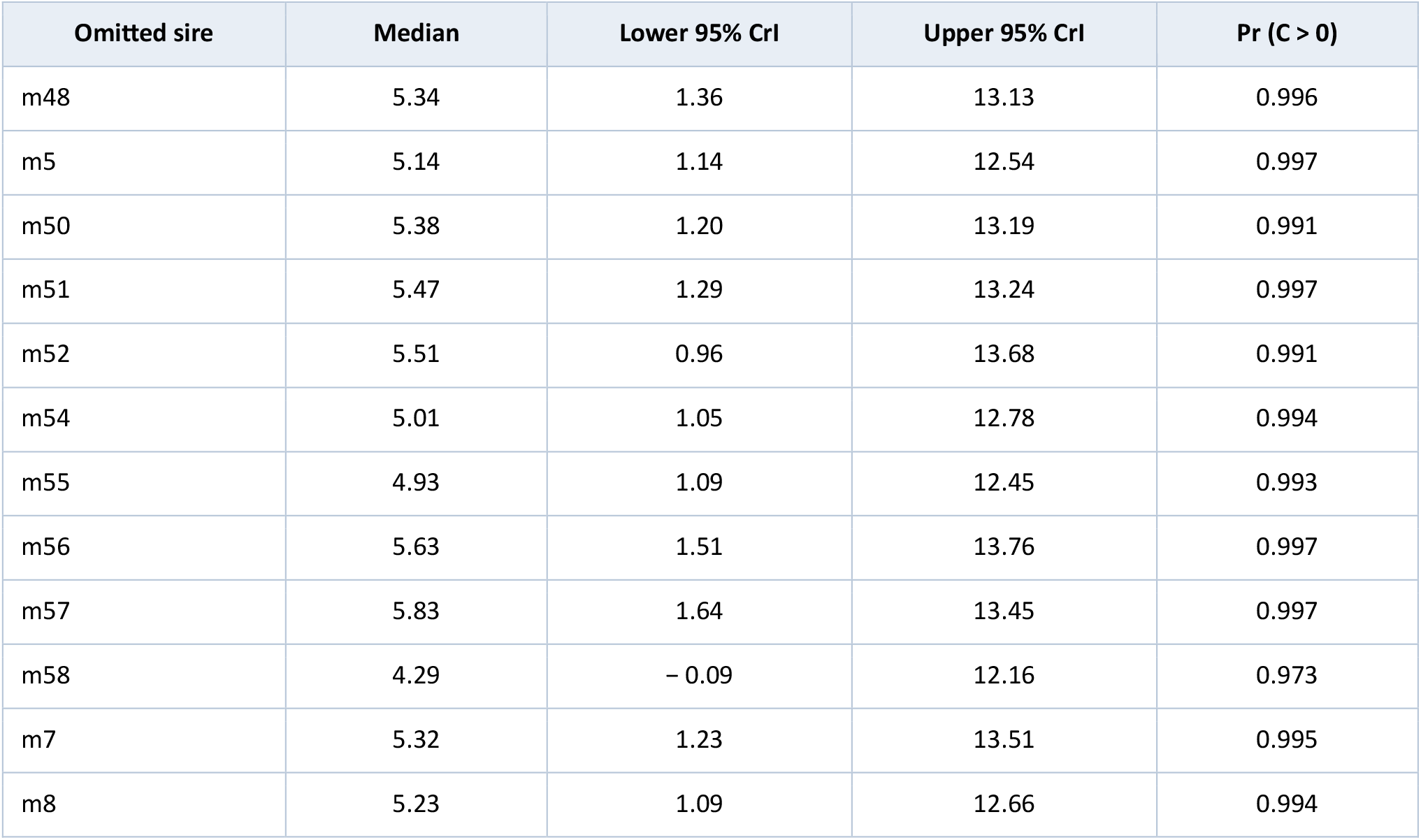
Leave-one-sire-out estimates of the between-trait log variance-ratio contrast for all 35 sires.

**Table S4.**
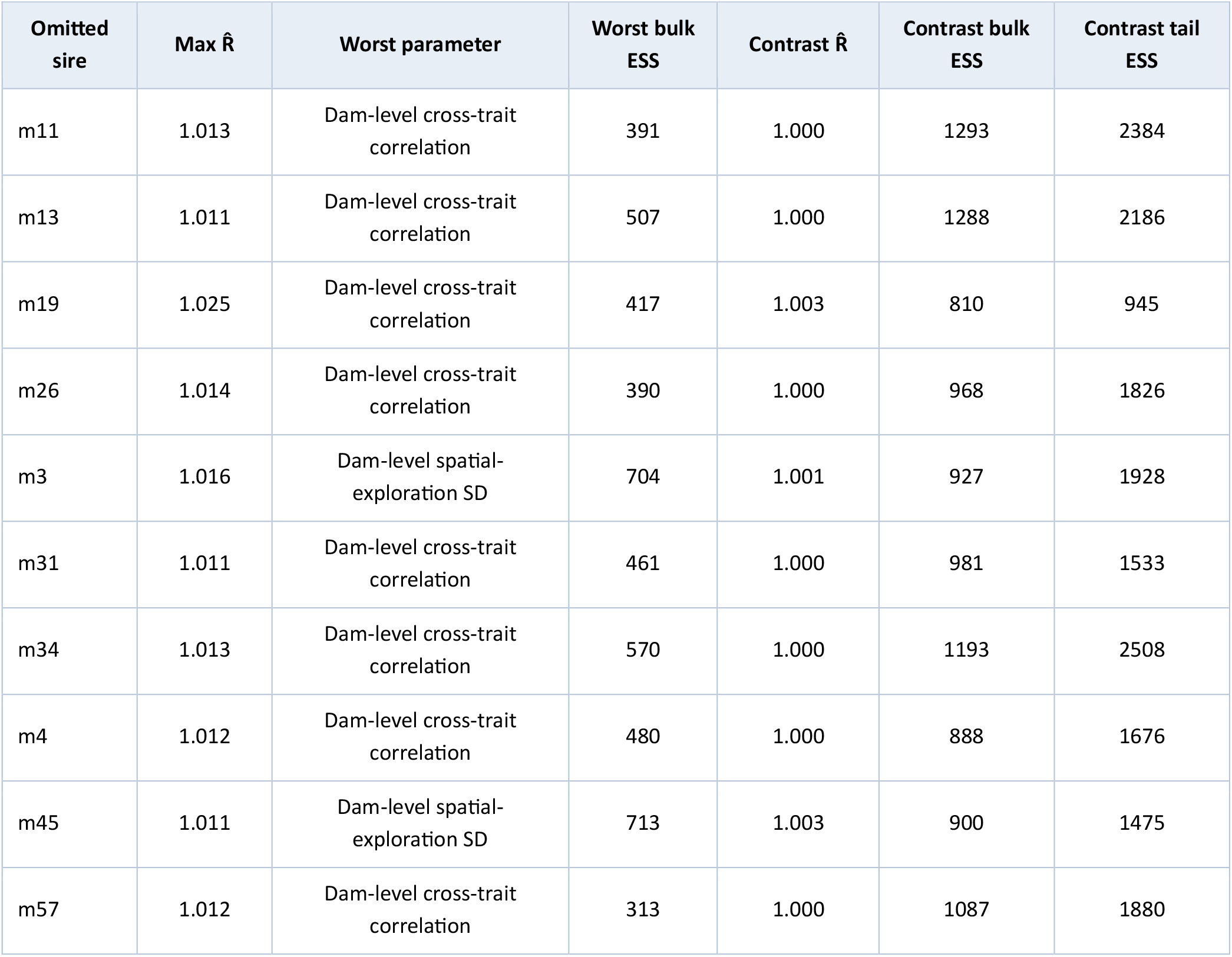
Targeted convergence audit for leave-one-sire-out models with a maximum parameter-level 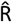 above 1.01.

**Figure S1.**
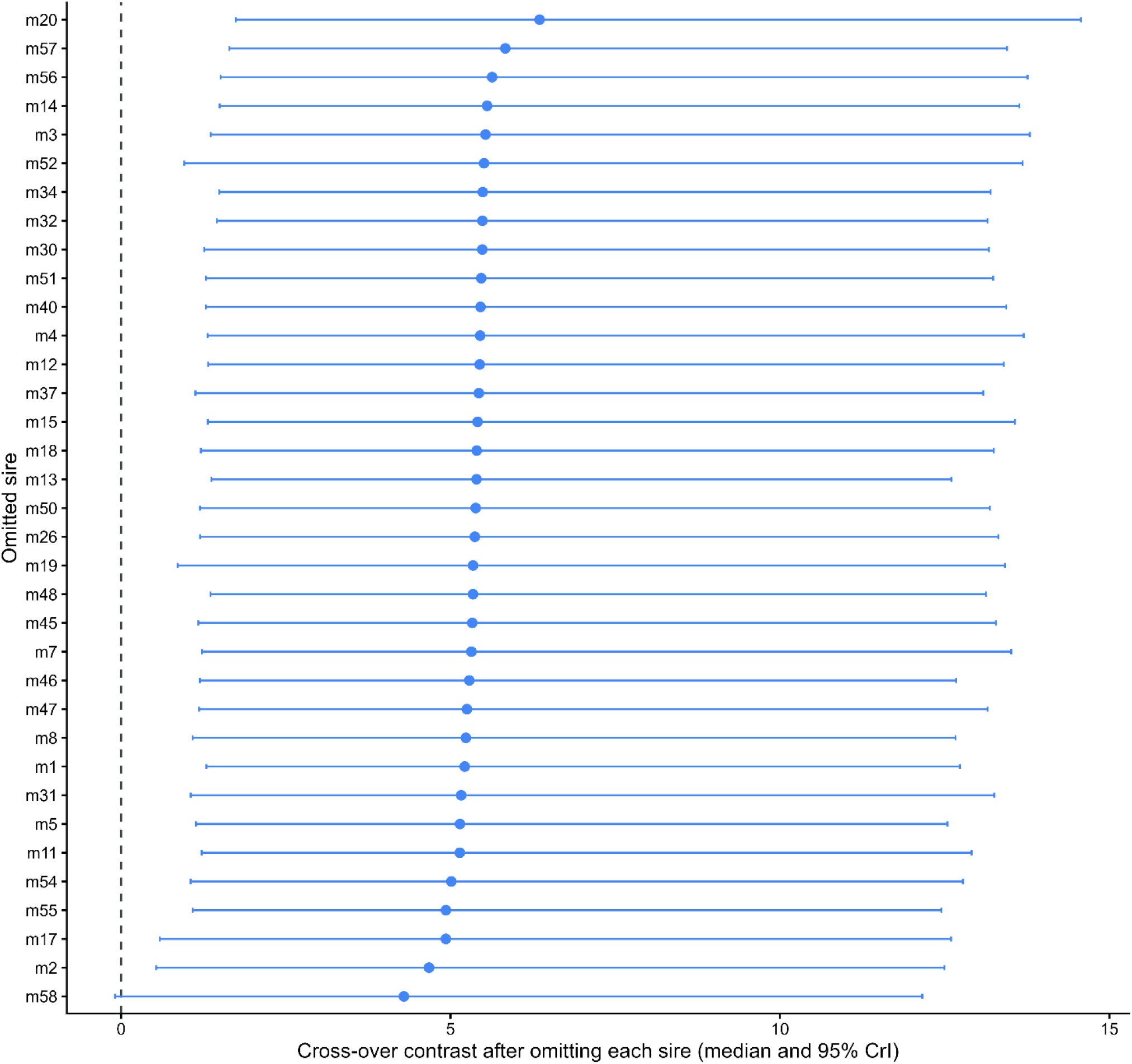
Leave-one-sire-out assessment of the between-trait log variance-ratio contrast. The primary bivariate model was refitted after excluding each of the 35 sires in turn. Points show posterior medians and horizontal bars show 95% credible intervals. The vertical dashed line denotes a contrast of zero. Positive values indicate a greater sire-to-dam variance ratio for exploration than for activity.

