## Supplementary Information for "Exploration and activity show contrasting familial resemblance"

### Model diagnostics and posterior predictive assessment

The primary bivariate model showed satisfactory sampling performance (Table S1). The maximum rank-normalised R̂ was 1.007, and the minimum bulk and tail effective sample sizes were 643 and 1111, respectively. No divergent transitions or maximum-tree-depth exceedances occurred.

Table S1. Sampling diagnostics for the primary and sensitivity models.

| **Model** | **Max R̂** | **Min bulk ESS** | **Min tail ESS** | **Divergences** | **Tree-depth hits** |
| --- | --- | --- | --- | --- | --- |
| Primary | 1.007 | 643 | 1111 | 0 | 0 |
| Weaker priors | 1.005 | 518 | 604 | 0 | 0 |
| Sires with ≥2 dams | 1.003 | 885 | 1041 | 0 | 0 |

### Sensitivity to prior specification and family structure

The between-trait log variance-ratio contrast was consistent under both alternative prior specification and when the analysis was restricted to sires represented by at least two dams (Table S2). Under weaker priors, the posterior median contrast was 5.54 (95% credible interval, CrI: 1.24 –13.30), and its posterior probability of being positive was 0.993. Restricting the analysis to sires represented by at least two dams produced a posterior median contrast of 5.14 (95% CrI: 0.86 – 12.73; posterior probability positive = 0.989). These estimates were similar to the primary model estimate of 5.32 (95% CrI: 1.14 – 13.32; posterior probability positive = 0.993). Sampling diagnostics were satisfactory for both sensitivity models (Table S1). Maximum rank-normalised R̂ values were 1.005 under weaker priors and 1.003 in the analysis restricted to sires with at least two dams. Neither model produced divergent transitions or maximum-tree-depth exceedances.

Table S2. Between-trait log variance-ratio contrast under the primary and sensitivity models. Positive values indicate a greater sire-to-dam variance ratio for exploration than for activity.

| **Model** | **Median** | **Lower 95% CrI** | **Upper 95% CrI** | **Pr (C > 0)** |
| --- | --- | --- | --- | --- |
| Primary | 5.32 | 1.14 | 13.32 | 0.993 |
| Weaker priors | 5.54 | 1.24 | 13.30 | 0.993 |
| Sires with ≥2 dams | 5.14 | 0.86 | 12.73 | 0.989 |

### Leave-one-sire-out influence analysis

Excluding each of the 35 sires in turn did not alter the direction of the between-trait log variance-ratio contrast (Figure S1; Table S3). Posterior median contrast estimates ranged from 4.29 to 6.35. The minimum posterior probability that the contrast was positive was 0.973. The most conservative estimate followed exclusion of sire m58 (posterior median = 4.29, 95% CrI: − 0.09 – 12.16; posterior probability positive = 0.973). Excluding sire m2 produced the next-lowest median estimate (4.67, 95% CrI: 0.53 – 12.50; posterior probability positive = 0.988). Thus, the positive between-trait contrast was not attributable to a single paternal family.

Ten reduced models had a maximum parameter-level R̂ marginally above 1.01 (Table S4). In eight cases, the maximum value occurred for the dam-within-sire-level correlation between exploration and activity, in the remaining two, it occurred for the dam-within-sire standard deviation for exploration. Convergence diagnostics for the derived contrast were satisfactory in all ten models: R̂ ranged from 1.000 to 1.003, bulk effective sample sizes from 810 to 1293, and tail effective sample sizes from 945 to 2508. No reduced model contained divergent transitions.

Table S3. Leave-one-sire-out estimates of the between-trait log variance-ratio contrast for all 35 sires.

| **Omitted sire** | **Median** | **Lower 95% CrI** | **Upper 95% CrI** | **Pr (C > 0)** |
| --- | --- | --- | --- | --- |
| m1 | 5.21 | 1.29 | 12.73 | 0.996 |
| m11 | 5.14 | 1.22 | 12.91 | 0.994 |
| m12 | 5.44 | 1.32 | 13.40 | 0.995 |
| m13 | 5.39 | 1.37 | 12.60 | 0.996 |
| m14 | 5.56 | 1.50 | 13.64 | 0.996 |
| m15 | 5.41 | 1.32 | 13.57 | 0.997 |
| m17 | 4.93 | 0.59 | 12.60 | 0.987 |
| m18 | 5.40 | 1.21 | 13.25 | 0.993 |
| m19 | 5.34 | 0.86 | 13.42 | 0.989 |
| m2 | 4.67 | 0.53 | 12.50 | 0.988 |
| m20 | 6.35 | 1.74 | 14.57 | 0.998 |
| m26 | 5.37 | 1.20 | 13.31 | 0.995 |
| m3 | 5.53 | 1.36 | 13.79 | 0.996 |
| m30 | 5.48 | 1.26 | 13.17 | 0.997 |
| m31 | 5.16 | 1.06 | 13.25 | 0.994 |
| m32 | 5.48 | 1.45 | 13.15 | 0.997 |
| m34 | 5.49 | 1.49 | 13.20 | 0.997 |
| m37 | 5.43 | 1.13 | 13.09 | 0.995 |
| m4 | 5.45 | 1.31 | 13.70 | 0.997 |
| m40 | 5.46 | 1.29 | 13.43 | 0.996 |
| m45 | 5.33 | 1.17 | 13.28 | 0.994 |
| m46 | 5.29 | 1.20 | 12.68 | 0.994 |
| m47 | 5.25 | 1.18 | 13.15 | 0.995 |
| m48 | 5.34 | 1.36 | 13.13 | 0.996 |
| m5 | 5.14 | 1.14 | 12.54 | 0.997 |
| m50 | 5.38 | 1.20 | 13.19 | 0.991 |
| m51 | 5.47 | 1.29 | 13.24 | 0.997 |
| m52 | 5.51 | 0.96 | 13.68 | 0.991 |
| m54 | 5.01 | 1.05 | 12.78 | 0.994 |
| m55 | 4.93 | 1.09 | 12.45 | 0.993 |
| m56 | 5.63 | 1.51 | 13.76 | 0.997 |
| m57 | 5.83 | 1.64 | 13.45 | 0.997 |
| m58 | 4.29 | − 0.09 | 12.16 | 0.973 |
| m7 | 5.32 | 1.23 | 13.51 | 0.995 |
| m8 | 5.23 | 1.09 | 12.66 | 0.994 |

Table S4. Targeted convergence audit for leave-one-sire-out models with a maximum parameter-level R̂ above 1.01.

| **Omitted sire** | **Max R̂** | **Worst parameter** | **Worst bulk ESS** | **Contrast R̂** | **Contrast bulk ESS** | **Contrast tail ESS** |
| --- | --- | --- | --- | --- | --- | --- |
| m11 | 1.013 | Dam-level cross-trait correlation | 391 | 1.000 | 1293 | 2384 |
| m13 | 1.011 | Dam-level cross-trait correlation | 507 | 1.000 | 1288 | 2186 |
| m19 | 1.025 | Dam-level cross-trait correlation | 417 | 1.003 | 810 | 945 |
| m26 | 1.014 | Dam-level cross-trait correlation | 390 | 1.000 | 968 | 1826 |
| m3 | 1.016 | Dam-level spatial-exploration SD | 704 | 1.001 | 927 | 1928 |
| m31 | 1.011 | Dam-level cross-trait correlation | 461 | 1.000 | 981 | 1533 |
| m34 | 1.013 | Dam-level cross-trait correlation | 570 | 1.000 | 1193 | 2508 |
| m4 | 1.012 | Dam-level cross-trait correlation | 480 | 1.000 | 888 | 1676 |
| m45 | 1.011 | Dam-level spatial-exploration SD | 713 | 1.003 | 900 | 1475 |
| m57 | 1.012 | Dam-level cross-trait correlation | 313 | 1.000 | 1087 | 1880 |

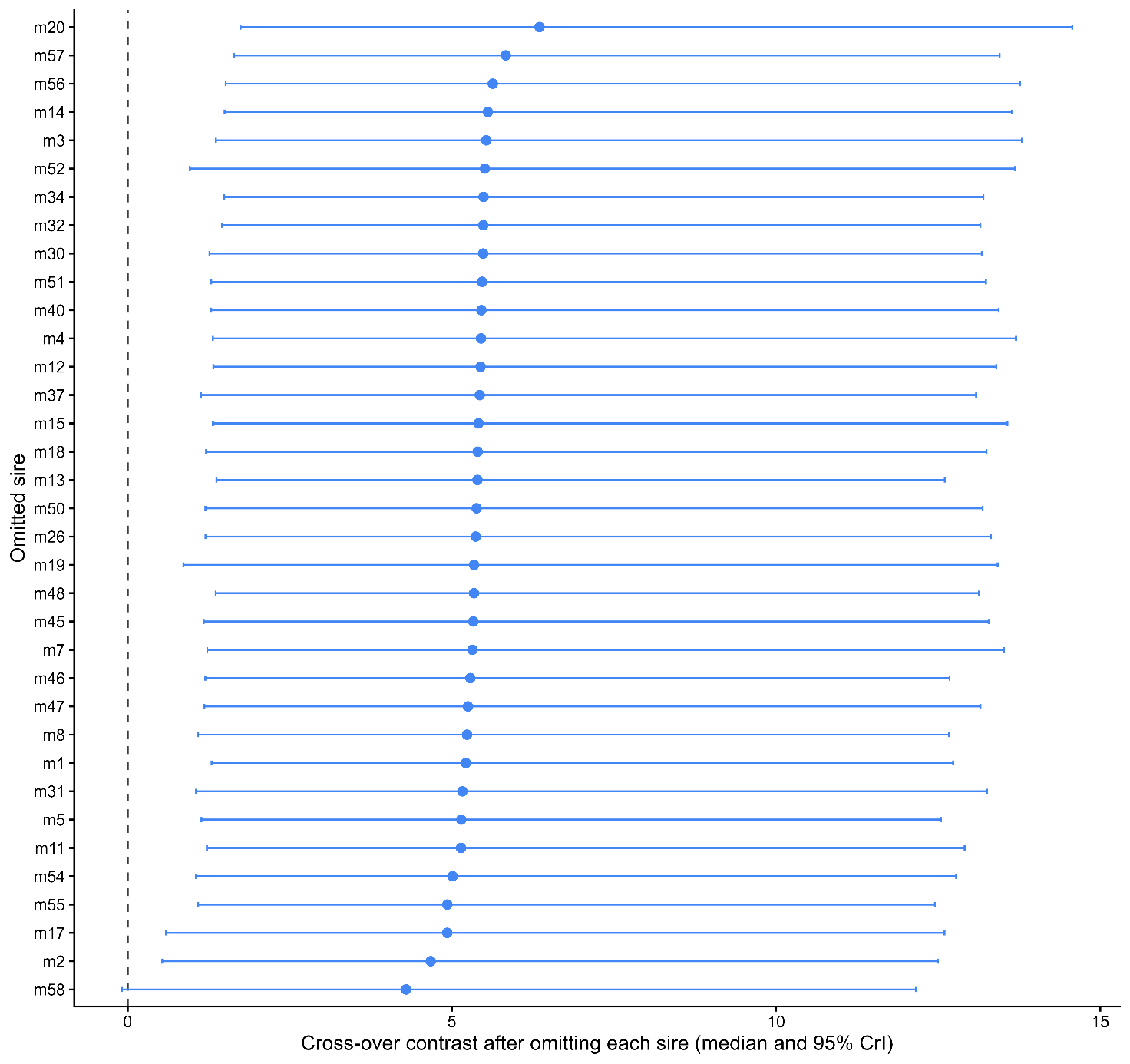

**Figure S1. Leave-one-sire-out assessment of the between-trait log variance-ratio contrast.** The primary bivariate model was refitted after excluding each of the 35 sires in turn. Points show posterior medians and horizontal bars show 95% credible intervals. The vertical dashed line denotes a contrast of zero. Positive values indicate a greater sire-to-dam variance ratio for exploration than for activity.
